# Neuromagnetic signatures of the spatiotemporal transformation for manual pointing

**DOI:** 10.1101/253328

**Authors:** G. Blohm, H. Alikhanian, W. Gaetz, H.C. Goltz, J.F.X. DeSouza, D.O. Cheyne, J.D. Crawford

## Abstract

Movement planning involves transforming the sensory goal representation into a command in motor coordinates. Surprisingly, the real-time dynamics of sensorimotor transformations at the whole brain level remain unknown, in part due to the spatiotemporal limitations of fMRI and neurophysiological recordings. Here, we used magnetoencephalography (MEG) during pro-/anti-wrist pointing to determine (1) the cortical areas involved in transforming visual signals into appropriate hand motor commands, and (2) how this transformation occurs in real time, both within and across the regions involved. We computed sensory, motor, and sensorimotor indices in 16 bilateral brain regions for direction coding based on hemispherically lateralized de/synchronization in the α (7-15Hz) and β (15-35Hz) bands. We found a visuomotor progression, from pure sensory codes in ‘early’ occipital-parietal areas, to a temporal transition from sensory to motor coding in the majority of parietal-frontal sensorimotor areas, to a pure motor code, in both the α and β bands. Further, the timing of these transformations revealed a top-down pro/anti cue influence that propagated ‘backwards’ from frontal through posterior cortical areas. These data directly demonstrate a progressive, real-time transformation both within and across the entire occipital-parietal-frontal network that follows specific rules of spatial distribution and temporal order.

## Introduction

Planning a goal-directed movement requires the transformation of visual signals into the motor signals suitable to activate the relevant muscle groups (Kalaska and Crammond 1992; Soechting and Flanders 1992; Andersen and Buneo 2002; Crawford et al. 2011). Distinguishing the spatial tuning of visual and motor signals can be challenging because stimulus and movement direction often correspond. One way to address this problem is by coupling neurophysiological or neuroimaging recordings with tasks that dissociate the stimulus from the response (Connolly et al. 2000; DeSouza 2002; Gail and Andersen 2006; Gail et al. 2009; Gertz and Fiehler 2015; Kuang et al. 2016; Cappadocia et al. 2017). However, the temporal limitations of fMRI do not allow recording of the real-time dynamics of sensorimotor transformations, whereas the local nature of invasive neurophysiological recordings do not allow observation of the sensorimotor transformations at the whole-brain level. Consequently, the dynamic, coordinated, mechanistic involvement of different human brain areas in visuomotor transformations, i.e. the distribution and order of cortical events in real time, remains largely unknown.

Previous neurophysiological, imaging, and neuropsychological studies suggest that the parietal-frontal network is responsible for the sensory-to-motor transformation underlying grasp (Michaels and Scherberger 2018) and reach (Buneo and Andersen 2006; Medendorp et al. 2011; Vesia and Crawford 2012) planning. It is generally believed that visual stimulus direction is compared to initial hand position to calculate a movement vector in a cortical network that includes superior parietal-occipital cortex (SPOC), mid-posterior intraparietal cortex (mIPS), and dorsal premotor cortex (PMd) (Pesaran et al. 2006, 2010; Khan et al. 2007; Chang et al. 2009; Vesia et al. 2010). Further, it is thought that in occipital-parietal cortex these parameters are coded relative to the eye, whereas they are transformed by the parietal-frontal network to result in effector-centered coordinates in frontal areas (Batista et al. 1999; DeSouza et al. 2000; Snyder 2000; Kakei et al. 2001, 2003; Fernandez-Ruiz et al. 2007; Khan et al. 2013; Fujiwara et al. 2017). Finally, it has recently been noted that these transformations are not entirely serial; it appears that human occipital cortex is reactivated, perhaps through reentrant pathways and perhaps through imagining the goal, during the planning and execution of movements (Singhal et al. 2013; Chen et al. 2014; Cappadocia et al. 2017).

Previous neuroimaging studies have capitalized on lateralized direction selectivity, i.e., right target/movement representation in left brain vs. left representation in right brain, to investigate visuomotor transformations in human cortex (Medendorp et al. 2003; Medendorp 2004; Beurze et al. 2007, 2009, 2010; Fernandez-Ruiz et al. 2007; Bernier et al. 2012; Vesia and Crawford 2012; Chen et al. 2014). In particular, this affords the opportunity to use simple stimulus-response dissociation tasks such as ‘anti-pointing’ to trace the progression of coding of remembered visual direction versus planned movement direction through a sensorimotor network (Curtis 2006). In anti-reaching (like anti-saccades), participants are instructed to point or reach in the opposite direction to the visual stimulus, sometimes after a delay. Such studies have generally found contralateral stimulus coding in occipital cortex during target representation and movement direction coding in parieto-frontal cortex during movement planning and execution (Connolly et al. 2000; Chen et al. 2014; Gertz and Fiehler 2015). Further, a slow event-related fMRI design has shown a progression from visual to motor coding both within and across areas in the occipital-parietal-frontal axis (Cappadocia et al. 2017). However, the neurovascular underpinnings of fMRI do not allow the real-time characterization of dynamics, e.g. the temporal ability to discriminate feed-forward from feed-back mechanisms, or clearly discriminate oscillatory network behavior from discrete spiking activity (Logothetis 2008; Kuang et al. 2016).

Conversely, single and multiunit recordings have the advantage of providing direct measures of neural activity and can discriminate action potentials from subthreshold and oscillatory network activity; e.g., Gail and colleagues have utilized this advantage in monkeys trained to perform the anti-reach task (Gail and Andersen 2006; Gail et al. 2009; Kuang et al. 2016). These experiments show that local field potentials in the primate parietal reach region (probably corresponding to mid posterior intraparietal and superior parietal-occipital cortex in the human) primarily encode the direction of visual input, whereas action potentials primarily encode the future movement direction. Like anti-saccade studies (Munoz and Everling 2004; Zhang and Barash 2004), these experiments show a capacity for multiple simultaneous codes and remapping of information within neurons. However, experiments in monkeys require extensive training to obtain results (potentially re-wiring the brain), are limited to one or a few brain areas, and may evoke species differences (Munoz and Everling 2004; Cappadocia et al. 2017).

While both fMRI and animal electrophysiology are critical and complementary techniques, there is also the need for a technique that might bridge the technical gap between them. Magnetoencephalography (MEG) is a promising candidate, because it provides simultaneous recordings from the entire brain in untrained humans, yielding a relatively direct link to ensemble neuronal activity (compared to the neurovascular coupling in fMRI). MEG also naturally provides frequency-dependent measures of brain oscillations, and millisecond temporal resolution. Spatial resolution and source localization in MEG have long been an issue, but this too has made recent advances (Alikhanian et al. 2013; Cheyne 2013). MEG has already been used to show that working memory-related γ band activity codes goal location rather than stimulus position in a delayed anti-saccade task (Van Der Werf et al. 2008). Such results point towards a gradual sensorimotor transformation across several brain areas, but a whole-brain analysis of anti-pointing has not yet been attempted.

Here, to reveal the dynamics and source of the sensory-to-motor transformation for manual control at the whole-brain level, we combined high spatial-temporal resolution MEG with a delayed pro-/anti-pointing task designed to dissociate sensory from motor activity. Based on current models of sensory-motor transformations (see Discussion for details), we predicted to see either a gradual feed-forward transformation from sensory to motor coding along cortical space, or a transformation from sensory to motor coding across time within a single (or small group of) area – presumably within parietal-premotor cortex. Isolating sensory, sensorimotor, and motor codes in real time using a whole-brain source reconstruction analysis, we show 1) a lateralized sensory-to-motor gradient along the occipital-to-parietal-to-frontal axis, 2) a progressive spatial transformation from sensory to motor coding both within and across these areas, and 3) an inverse frontal-to-posterior temporal transformation in response to the top-down pro/anti instruction.

## Methods

We used MEG to investigate the visual-to-spatial reference frame transformation in human cortex. MEG signals result through Maxwell’s law from the net local dendritic ionic currents produced during synaptic transmission in pyramidal cell layers of cortex (Murakami and Okada 2006; Koser 2010; Lopes da Silva 2013). MEG surface signals can be used to perform source reconstruction which is largely immune to tissue boundary effects (unlike EEG), resulting in high temporal AND spatial resolution (Baillet 2017). Since MEG signals are most detectable from currents that are tangential to the scalp, the most reliable signals can be measured from cortical sulci (Hillebrand and Barnes 2002).

Natural dynamics result in a variety of emergent, endogenous rhythms in the central nervous system. Synchronized oscillations in neural ensembles can occur due to synchronization from oscillatory activity of distant connected brain areas (Pikovsky et al. 2003), or due to intrinsic neuronal properties and local connectivity leading to synchronous population resonance effects (Zeitler et al. 2009). Amplitude changes in ongoing natural rhythms (e.g., α: 7-15 Hz; and β: 15-35 Hz) can result from changes in local de/synchronization induced by sensory-motor or other events (Salmelin and Hari 1994; Pfurtscheller et al. 1998; Pfurtscheller and Lopes da Silva 1999; Neuper and Pfurtscheller 2001; Jurkiewicz et al. 2006). We take advantage of this coupling to investigate how different cortical areas process sensory and motor information in a differential manner (Koser 2010).

### Participants

We recorded data from ten healthy adult participants (eight males and two females, 22-45 years old). We used this number of participants because we can collect many more repetitions per participant in MEG experiments than in typical fMRI experiments, and due to the mathematically precise nature of electro-magnetic coupling, appropriate statistics reduce the risk of false statistical positives (see Statistical Methods). Participants were screened prior to participation in this study; none of the participants had any known history of neurological dysfunction, injury or metallic implants and all (but one with amblyopia) participants reported normal or corrected-to-normal vision. This study was approved both by the York University and The Hospital for Sick Children research ethics boards. All participants gave informed consent.

### MEG setup, behavioral recordings and anatomical MRIs

Participants sat upright in a magnetically shielded room (Vacuumschmelze Ak3b) with their head in the dewar (see Figure 1C) of a whole-head 151-channel (axial gradiometers, 5 cm baseline) CTF MEG system (VSM Medtech, Coquitlam, Canada) at The Hospital for Sick Children. Noise levels were below 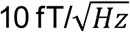 above 1.0 Hz. Prior to MEG data acquisition, each participant was fitted with coils placed at three fiducial landmarks (nasion and pre-auricular points) that were localized by the MEG acquisition hardware to establish the position of the participant’s head relative to the MEG sensors at the beginning and end of each recording. Coil placements were carefully measured and photographed for off-line co-registration of recorded MEG data to anatomical MR images obtained for each participant.

**Figure 1:**
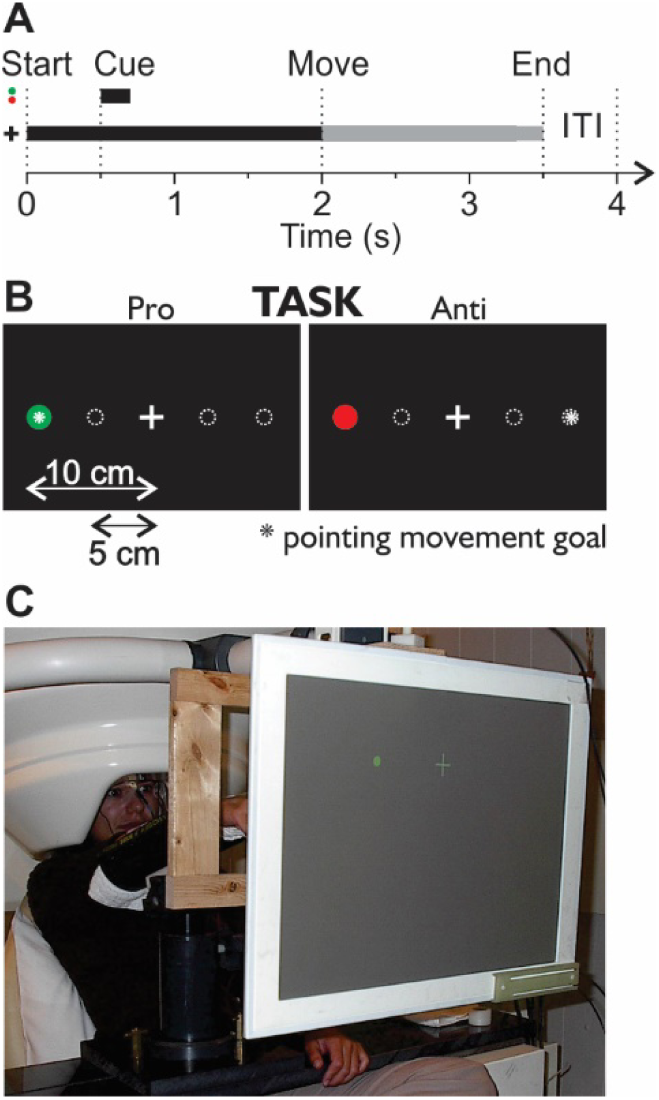
Set-up. **A**. Time line of trials. Each trial started with a fixation cross. 500ms later, the cue was flashed for 200ms at one of 4 possible locations (see panel B). The cue was either red or green indicating whether subjects were in a pro- or anti-condition. After the cue onset, there was a 1500ms memory delay. Then the fixation cross was dimmed, which was the movement instruction for the participants. Participants had to perform either a pro- or an anti-pointing movement and had 1500ms to do so. Then the fixation cross disappeared for 500ms to indicate the end of the trial. Subjects were instructed to return to the central initial hand position during this period. **B**. Visual arrangement of display. Subjects were instructed to keep fixating the white cross during the whole trial. The gray doted outlines indicate the potential locations of the cue. The screen was approximately 80cm distant from the subjects and targets were located at 5 and 10cm on either side of the fixation cross. **C**. Photograph of the setup. Subjects sat upright in the MEG apparatus with their head inside the dewar. Their arm was held by a forearm rest. We used the wooden frame to hold the light interrupters that detected when subjects pointed to the left or right.

We measured participants’ horizontal eye movements through electro-oculography (EOG) using two bipolar temporal electrodes. In order to measure horizontal wrist pointing movements (see ‘Task’ below), we also measured surface electromyographic (EMG) activity from placements on the lateral and medial aspects of the anterior and posterior forearm with four pairs of 1 cm diameter electrodes to measure both wrist flexion / extension and radial / ulnar deviation during different hand postures. Those pairs were placed over the Extensor Carpi Radialis Longior (ECRL), Extensor Communis Digitorum (ECD), Extensor Carpi Ulnaris (ECU), and Supinator Longus (SL) muscles. Electrodes for both EMG and EOG recordings were Ag/AgCl solid gel Neuroline (Ambu) electrodes of type 715 12-U/C. EMG and EOG channels were collected as auxiliary channels of the CTF MEG recording system at the same on-line filter settings and sample rate. In order to simplify EMG-based movement detection and movement direction, we also recorded the time when the participant’s finger passed through one of two LED light paths mounted about 2cm left and right of straight-ahead wrist position.

Visual stimuli were back-projected onto a translucent tangential screen at a distance of 1-m. Stimuli were rear-projected through the shielded room wall using a LDC projector (Sanyo PLC-XP51) with a zoom lens (Navitar model 829MCZ087) and controlled in real time by the Presentation program (Neurobehavioural Systems, Inc., Albany, CA, USA) with a refresh rate 60 Hz. Timing and condition information for each trial was sent to the CTF MEG recording system through a parallel port cable and was recorded in real time.

We also obtained structural (*T*_1_-weighted, 3D-SPGR) MRI scans from a 1.5 T Signa Advantage System (GE Medical Systems, Milwaukee, WI) on the same day. These scans were used for co-registration of the MEG dewar-based coordinate system to each participant’s brain coordinates by identifying the locations of the head localization coils on orthogonal slices of each participant’s MRI. For each participant, the inner skull surface was derived from *T*_1_-weighted MR data using the BrainSuite software package (Shattuck and Leahy 2002).

### Task

To dissociate between sensory and motor coding in the brain, we designed a pro-/anti-pointing task (Figure 1A-B). At the beginning of each trial, a white fixation cross appeared at the center of the screen at eye level and participants were required to fixate that cross throughout the trial. 500 ms later, a green or red dot (5 mm diameter) appeared for 200ms either 5 cm (near target) or 10 cm (far target) left or right of the fixation cross. Target color indicated whether participants had to point towards the target (green: pro trials) or to its mirror opposite location (red: anti trials). Target color codes were counter-balanced across participants. Presenting the instruction cue together with the goal was crucial to our experimental design to uncover the sensory-to-motor transformation in real time. After a 1,500ms delay, the movement was cued by dimming the fixation cross on the screen. Participants were asked to point as accurately as possible by producing wrist movements that matched the goal distance and direction. Participants received performance feedback from the experimenter during practice trials prior to the experimental recordings.

Participants performed four blocks of 400 trials each of this task. Each block contained a pseudo-random balanced set of 50 trials for each condition (combination of: close/far, left/right, pro/anti). They carried out three blocks with their right arm extended and right hand in the pronation (palm facing down), upright (palm facing leftward) and down (palm facing rightward) postures respectively. We included two different target eccentricities in each hemifield to encourage participants to program each individual movement instead of recalling a default left or right movement, but we averaged across both eccentricities for left and right conditions in the data analysis.

### Data processing

All analyses were done in Matlab (The Mathworks, Inc., Natick, MA, USA). MEG, EMG and EOG data were collected at 625 Hz. MEG data were online low-pass filtered at 200 Hz using synthetic third-order gradiometer noise cancelation. For detection of pointing movement onset, EMG data were band-pass filtered offline from 15 Hz to 200Hz and full-wave rectified. Movement onsets were marked using an automated algorithm that detected when the EMG signals rose above 3 times the standard deviation of the baseline EMG activity (measured before target onset). The first detection across all four muscles was used as the movement onset time and visually inspected and manually corrected, if necessary (<2% of trials). All data were then aligned to both cue onset (-500 ms to 1,500 ms around cue onset) and movement onset (-1,500 ms to 500 ms around movement onset) and extracted for further analysis. All data were visually inspected and trials with movement direction errors were discarded from further analysis (3.2% of total trials across participants).

We reconstructed instantaneous source power from the raw MEG sensor data using a scalar, unit noise gain minimum-variance beamforming algorithm (Cheyne et al. 2007). This inverse method has been shown to achieve high localization accuracy under conditions of low to moderate SNR (Sekihara et al. 2005; Neugebauer et al. 2017). All further analyses were conducted in source space. For spatial averaging across participants, individual participants’ source activity was transformed into MNI coordinate space using standard affine transformations (linear and non-linear warping) in SPM 8 and then projected onto a surface mesh of an average brain (PALS-B12 atlas (Van Essen 2005)) using Caret (Van Essen et al. 2001). To identify consistently activated brain regions, we used an adaptive clustering approach (Alikhanian et al. 2013). Note that projected images were for visualization only and the identification of brain areas operates on the raw, non-contrasted, time-averaged whole-brain activations and was thus orthogonal to our stimulus-condition contrasted analysis (see below), making this approach statistically valid and sound (Kriegeskorte et al. 2009; Kilner 2013). We then estimated source time courses (“virtual sensors”) at the identified areas and extracted trial-to-trial source activity at those locations (using the same event-related beamformer algorithm to estimate the spatial filter weights). This source data was used to compute time-frequency responses (TFRs) using the Brainwave Matlab toolbox developed at the Hospital for Sick Children (http://cheynelab.utoronto.ca/brainwave) and custom code.

In all analyses, we considered the 500ms window prior to cue onset (-500 to 0ms) as the baseline (see Figure 1). All signals were referenced (i.e. normalized with respect to) to the frequency-dependent average power of the baseline period. For average whole-brain analyses, individual participant source power was estimated for a given frequency range using the above-mentioned beamformer and individually referenced before averaging. Whole-brain activity plots were thresholded based on signal power, not statistical significance to better appreciate source power. Data from left and right hemispheres were subtracted from one another to collapse data across corresponding bilateral brain areas, as this is commonly done (e.g. (Van Der Werf et al. 2008)). Similarly, TFR analyses were computed at the individual participant level as the relative changes of oscillatory power with respect to baseline and then averaged across participants.

### Data analysis

We carried out an event-related analysis of MEG data during our delayed pro-/anti-pointing task. To do so, we aligned individual trials to either cue onset times or wrist movement onset times, as measured by EMG. A typical trial is shown in Figure 2. Time series for EOG, EMG and selected MEG channels are shown along with selected subsets of parieto-occipital MEG channels. Movement onset for this particular trial can be seen easily in the EMG signals after the movement cue and EOG shows good fixation performance. Individual participant movement times are summarized in Table 1. In our subsequent analysis, we focused on the conventional α (visual) and β (motor) frequency bands of MEG signals, because these frequency bands have been previously associated with sensorimotor system activity in a variety of motor tasks (Neuper and Pfurtscheller 2001; Neuper et al. 2006; Koser 2010; Cheyne 2013; Lopes da Silva 2013; Spitzer and Haegens 2017).

**Table 1:** Movement time analysis. Mean, STD and percentiles are shown in s after cue onset. The move instruction was at 1.5s.

| *Subj. ID* | *Mean* | *STD* | *10%* | *25%* | *50%* | *75%* | *90%* |
| --- | --- | --- | --- | --- | --- | --- | --- |
| 1 | 1.9412 | 0.1530 | 1.7904 | 1.8464 | 1.9184 | 2.0096 | 2.1104 |
| 2 | 1.8547 | 0.1295 | 1.7248 | 1.7760 | 1.8416 | 1.9168 | 2.0078 |
| 3 | 1.7856 | 0.2198 | 1.6371 | 1.6912 | 1.7552 | 1.8468 | 1.9408 |
| 4 | 1.9606 | 0.4440 | 1.6128 | 1.7440 | 1.8960 | 2.1920 | 2.5360 |
| 5 | 1.6868 | 0.1783 | 1.5344 | 1.6176 | 1.6912 | 1.7492 | 1.8128 |
| 6 | 1.6881 | 0.2639 | 1.4384 | 1.6064 | 1.6976 | 1.7952 | 1.9120 |
| 7 | 1.7593 | 0.1239 | 1.6704 | 1.7056 | 1.7488 | 1.8000 | 1.8528 |
| 8 | 1.7565 | 0.4443 | 1.3344 | 1.5552 | 1.6800 | 1.9440 | 2.3232 |
| 9 | 1.7696 | 0.1888 | 1.6496 | 1.6928 | 1.7456 | 1.8064 | 1.8928 |
| 10 | 1.6031 | 0.1404 | 1.4176 | 1.5232 | 1.6256 | 1.6752 | 1.7328 |
| Merged | 1.7890 | 0.2786 | 1.5680 | 1.6704 | 1.7600 | 1.8768 | 2.0416 |
| Averaged | 1.8154 | 0.2642 | 1.5937 | 1.6893 | 1.7749 | 1.9048 | 2.0826 |
| ±STD | ±0.0829 | ±0.1090 | ±0.1330 | ±0.0855 | ±0.0833 | ±0.1261 | ±0.2034 |

**Figure 2:**
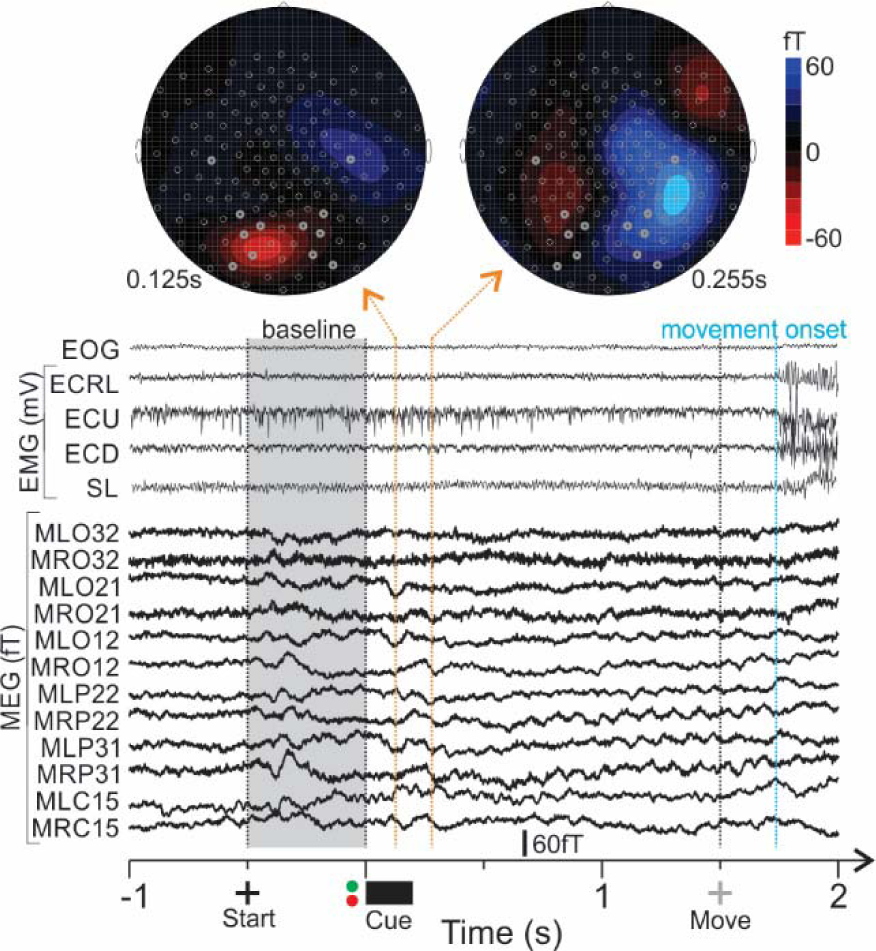
Typical trial. Time series of EOG, EMG and example MEG channels are shown for a single trial. Snapshots of scalp potentials at two different time points (125ms and 255ms after cue onset) are represented above the time series and show how the MEG amplitude can change over scalp space and time within a single trial. Flat EOG signal shows good fixation performance. EMG signals clearly demonstrate movement onset in Extensor Carpi Radialis Longior (ECRL), Extensor Communis Digitorum (ECD), Extensor Carpi Ulnaris (ECU), and Supinator Longus (SL) muscles. MEG channel labels (starting with M) indicate right/left occipital lobe (LO and RO), right/left parietal lobe (RP and LP) and right/left central lobe (LC and RC) sensor locations (bold sensors on scalp potential plots). We used the 500ms fixation period prior to cue onset as the baseline for all further analyses.

Our behavioral paradigm was designed to dissociate sensory from motor-related activity. In particular, pro- and anti-pointing trials required the same spatial stimulus to be transformed into different motor plans. In addition, we made use of the brain’s lateralization in spatial coding to separate sensory and motor coding in a fashion similar to the analyses employed successfully in recent neurophysiology (Kuang et al. 2016) and neuroimaging (Gertz and Fiehler 2015; Cappadocia et al. 2017) anti-reach studies. We thus extracted visual-spatially and motor-spatially selective brain activation by subtracting TFR results obtained from right and left stimulus/movement direction (depending on the analysis). Specifically, we hypothesized that ***sensory coding*** should discriminate between left and right stimulus location and should be independent of motor outcome (pro or anti). How this works is illustrated in Figure 3. To look for sensory coding, we thus computed the differential TFR of the following experimental conditions:

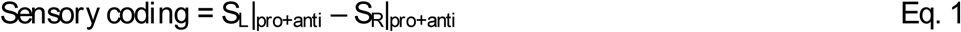

where S_L/R_ stands for left and right stimulus location irrespective of movement direction, i.e. adding pro- and anti-effects together. By adding pro- and anti-trials together, any motor-related effects should average out because pro- and anti-trials result in spatially oppositely directed movements. This is true even if there were posture-holding related muscle activity because such activity would also be present in the baseline period (subtracted). In addition, we tried to minimize such muscle activity by using a forearm rest (see Fig. 1C). In other words, a lateralized motor effect would produce opposite TFR results for pro- and anti-trials (de- vs re-synchronization), which – added together – will cancel out in the sensory coding computation. Any significant component in the TFR would thus point towards a sensory code in the brain area under investigation. Conversely, to look for ***motor coding***, we subtracted pro- and anti-trials because both result in opposite movements, thus emphasizing this difference while subtracting out any sensory coding effects, as illustrated in Supplementary Figure 1:

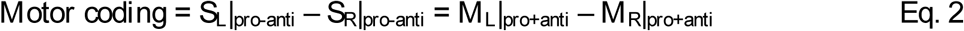

where M_L/R_ stands for left and right movement directions irrespective of stimulus location. This differential activity for investigating sensory coding and motor coding was computed both for the sensory (cue) and motor (movement) alignments of the data, i.e. highlighting early (stimulus-related) and late (movement-related) coding schemes during the sensory-to-motor transformation. (Note, that we can also compute a rule coding index to look for pro vs anti-task coding; however, this was beyond the purpose of this paper.)

**Figure 3:**
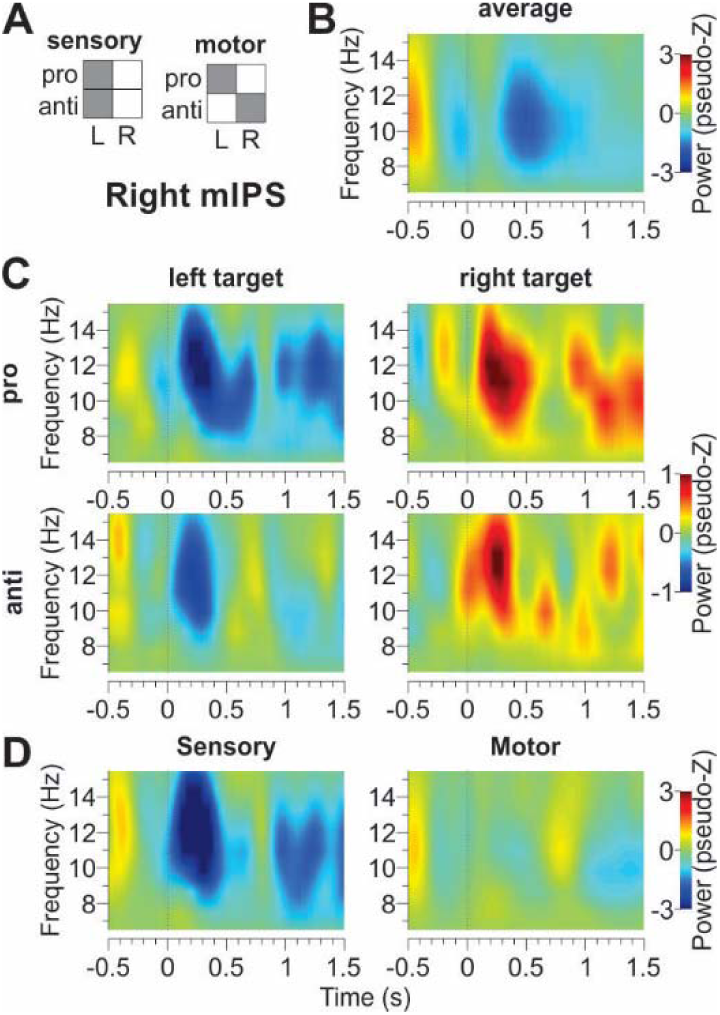
Sensory and motor coding predictions. **A**. Prediction for time-frequency response (TFR) results if an area codes sensory information (left) or motor information (right). For sensory information, activation patterns should be similar for left (L) and right (R) target locations, irrespective of the actual movement required, i.e. whether it is a pro- or anti-trial. For motor information, activation patterns should be similar for left (proL, antiR) and right (antiL, proR) movement directions, irrespective of the cue location. **B**. Stimulus-related averaged α-band TFR of the right medial intraparietal sulcus (mIPS) aligned to cue onset and averaged across pro, anti, L and R (red: re-synchronization; blue: desynchronization). **C**. Average-subtracted TFR for individual conditions for right mIPS. The similarity of left pro and anti, and right pro and anti TFRs points towards a sensory code. Since visual stimuli result in an overall bilateral de-synchronization in the visual system and we’re only interested in the differences between conditions, we subtracted the average across all conditions for presentation purpose (this was not done for our analyses, such as panel D). **D**. Subtracted activations for sensory coding (left panel, eq. 1) and motor coding (right panel, eq. 2).

Despite showing time intervals before cue onset and after movement onset, we only used the delay period activity between cue and movement to interpret our findings (not activity after movement onset). In this analysis we were not interested in posture effects; thus, we averaged data across all three arm postures to increase the number of trials / condition. While this might attenuate muscle-related MEG activity due to posture tuning effects, our results show consistent motor effects regardless.

We also computed a sensory-motor coding index to capture whether a given bilateral brain area predominantly codes information in sensory or motor coordinates. To do so, we used the above sensory coding (SC) and motor coding (MC) schemes as follows:

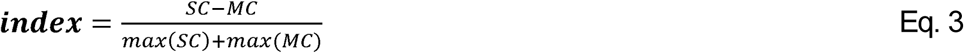

The sign of SC and MC were adjusted so that all main effects were positive. As a result, index=1 corresponds to perfect sensory coding, whereas index=-1 would be ideal motor coding (note that index=±1 values can only be obtained in noise-free data). We used the dominant α-band and β-band frequencies (10Hz and 20Hz, 1Hz band width) respectively and computed this index separately for each frequency and each trial, resulting in mean±SD for each time point and frequency. We then combined both time series in a statistically optimal fashion independently at each time point according to standard unbiased Bayesian integration with a Gaussian assumption. Results in Figure 8 are thus across-frequency sensory-motor coding indices.

### Statistical analysis

All contrasts (Eqs. 1-3) were computed for a given ROI source location from the clustering results on an individual-participant level and then averaged across participants. Statistical significance tests were conducted at the participant population level for individual time series of a given frequency based on TFR analyses. To evaluate statistical significance of this single-source analysis, we determined when the source power / sensory-motor index across participants was different from zero for at least 100ms (temporal clustering) in order to account for the multiple comparison problem (Maris and Oostenveld 2007). We used a conservative combined criterion, i.e. both 2-sided t-test and Wilcoxon rank-sum tests had to be significant (p<0.05) for a given time point. Since we identified significantly activated relevant brain areas using adaptive clustering (Alikhanian et al. 2013), thus did not perform statistical testing for whole-brain plots.

## Results

### General predictions and results

Since we were interested in the dynamics of individual brain areas during the sensory-to-motor transformation, we first performed source reconstruction and identified relevant brain areas showing significant activation in our task for each individual participant using adaptive clustering (see Methods). A total of 16 identified brain areas of interest were detected in each hemisphere and the coordinates are summarized in Table 2 for both hemispheres. We then placed virtual sensors in these locations and extracted single-trial time courses for each participant. This allowed us to compute time-frequency-responses (TFRs) separately for each participant and condition and analyze results across participants to obtain between-participant statistics. Note that (unlike BOLD activation in fMRI), the relevant variables here are desynchronization and resynchronization, both indicating a change in functional processing. Resynchronization is believed to arise from an increase of synchronous spike timing or membrane fluctuations in neurons and generally arises from internal recurrent processing in the brain, such as observed during motor preparation; desynchronization is observed when the natural rhythm of a brain area is disrupted, as is the case when sensory signals are being processed (Haken 1996; Pfurtscheller and Lopes da Silva 1999; Koser 2010).

**Table 2:**
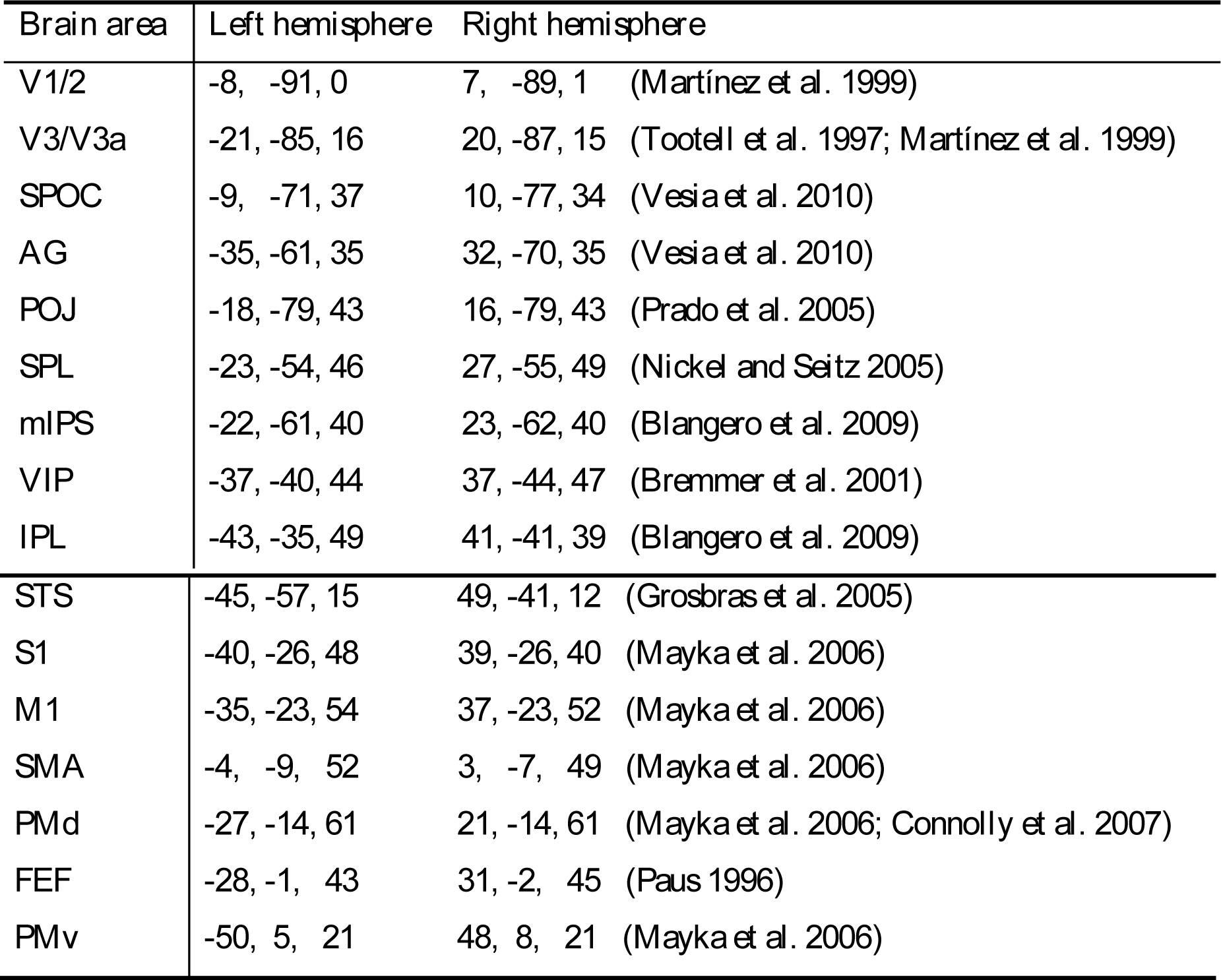
Average Talairach coordinates (mm) of functional brain areas. Areas were identified using an adaptive clustering approach (Alikhanian et al. 2013) and cross-validated from literature (indicated by references).

| ***Brain area*** | ***Left hemisphere*** | ***Right hemisphere*** |
| --- | --- | --- |
| V1/2 | -8, -91, 0 | 7, -89, 1 (Martínez et al. 1999) |
| V3/V3a | -21, -85, 16 | 20, -87, 15 (Tootell et al. 1997; Martínez et al. 1999) |
| SPOC | -9, -71, 37 | 10, -77, 34 (Vesia et al. 2010) |
| AG | -35, -61, 35 | 32, -70, 35 (Vesia et al. 2010) |
| POJ | -18, -79, 43 | 16, -79, 43 (Prado et al. 2005) |
| SPL | -23, -54, 46 | 27, -55, 49 (Nickel and Seitz 2005) |
| mIPS | -22, -61, 40 | 23, -62, 40 (Blangero et al. 2009) |
| VIP | -37, -40, 44 | 37, -44, 47 (Bremmer et al. 2001) |
| IPL | -43, -35, 49 | 41, -41, 39 (Blangero et al. 2009) |

|  |  |  |  |
| --- | --- | --- | --- |
| STS | -45, -57, 15 | 49, -41, 12 | (Grosbras et al. 2005) |
| S1 | -40, -26, 48 | 39, -26, 40 | (Mayka et al. 2006) |
| M1 | -35, -23, 54 | 37, -23, 52 | (Mayka et al. 2006) |
| SMA | -4, -9, 52 | 3, -7, 49 | (Mayka et al. 2006) |
| PMd | -27, -14, 61 | 21, -14, 61 | (Mayka et al. 2006; Connolly et al. 2007) |
| FEF | -28, -1, 43 | 31, -2, 45 | (Paus 1996) |
| PMv | -50, 5, 21 | 48, 8, 21 | (Mayka et al. 2006) |

To investigate whether a brain area would carry sensory or motor information (or both), we extracted the sensory code and motor code from our data, as described in the methods. The underlying assumption was that if a brain area only involved sensory aspects, then only the target location would determine the activation of that area, independently of the movement direction, i.e. regardless of whether it was a pro- or anti-trial. Conversely, motor coding would result in same TFR activation patterns for same movements, independently of target location.

Our process for computing sensory vs. motor coding is illustrated in Fig. 3 using Right mIPS and Supplementary Figure 1 using Left PMv as examples (see Fig. 4 legend for area acronym definitions). Panel A shows the idealized predictions for sensory and motor coding respectively. For sensory coding the responses to left (or right) targets should be similar, regardless of the pro-anti instruction. In the case of motor coding, the ‘anti’ instruction reverses the movement direction, so now the diagonals should provide matching data. Panels B shows the average TFR across left/right targets and pro/anti conditions, which we use as a baseline to subtract directionally non-specific activation. Panels C shows the average-subtracted activation for each condition separately, spatially arranged in the same fashion as the prediction panels. In the case of mIPS the resulting pattern clearly resembles the sensory prediction (i.e., red/resynchronized in right panels and blue/desynchronization in left panels). Other areas showed a pattern more consistent with motor coding (e.g., PMv, shown in Supplementary Figure 1). Finally, the data were subtracted (so that either the sensory response sums (and motor cancels) or vice versa, as shown in panel D of Figure 3 (and Supplementary Figure 1). As expected, this results in an early, predominantly sensory code for mIPS (Figure 3) and a late, predominantly motor code for PMv (Supplementary Figure 1). For the remainder of our analysis, we used these sensory coding and motor coding subtractions to represent our results.

**Figure 4:**
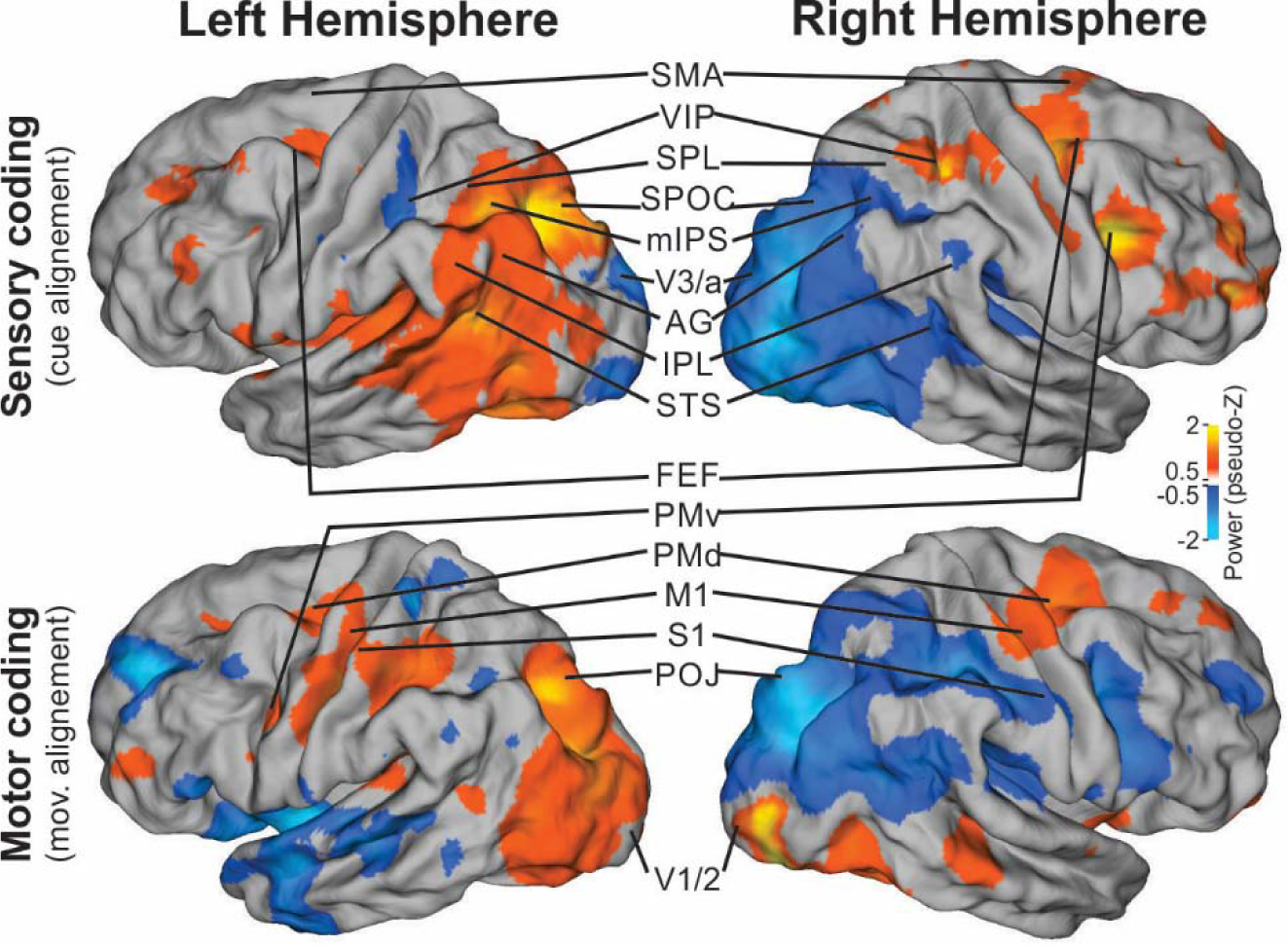
Whole-brain identified sensory and motor areas. Average 7-35Hz source power across all participants for sensory (top, cue-aligned, 0-500ms after cue onset are averaged) and motor (bottom, movement aligned, -500-0ms before movement onset are averaged) coding separately. Analyzed brain areas are highlighted. SMA: supplementary motor area; VIP: ventral intraparietal are; SPL: superior parietal lobe; mIPS: medial intraparietal sulcus; SPOC: superior parietal-occipital cortex; V3/a: visual area 3/3a; AG: angular gyrus; IPL: inferior parietal lobe; STS: superior temporal sulcus; FEF: frontal eye fields; PMv: ventral premotor area; PMd: dorsal premotor area; M1: primary motor cortex; S1: primary somato-sensory cortex; POJ: parietal-occipital junction; V1/2: primary visual areas 1/2.

To provide an overview of the results of this analysis, we performed these sensory and motor subtractions, averaging across both time (500ms window) and frequency (7-35 Hz) at the whole-brain level, averaged across participants, and then rendered the results over an average brain (see Methods). This is illustrated in Figure 4, which shows sensory coding over the period of 0-500ms following cue onset (upper panels) and motor coding during the period of -500 to 0ms preceding movement onset (lower panels), for both cortical hemispheres. On average, these datasets were separated by 815±83ms (see Table 1). The convention for this rendering is based on presentation of leftward targets/movements, where (due to the above-detailed subtractions), blue represents desynchronization and red represents resynchronization of activity relative to baseline. We also highlight average locations of the brain areas identified for each participant for further analysis (Table 2).

From this first pass, several trends emerge. First, for sensory coding during the cue response (Figure 4 top panels) a massive patch of occipital-temporal-parietal cortex containing V3, SOC, mIPS, SLP, AG, IPS, and STS shows sensory coding in response to the cue, with contralateral desynchronization and ipsilateral resynchronization (exceptions being the opposite trend in some posterior areas of left cortex and several frontal areas in right cortex). Second, for motor tuning preceding the action (lower panels) these areas show a somewhat diminished extent of contralateral desynchronization / ipsilateral resynchronization in their motor tuning, but this trend also appears in more frontal somatomotor areas like S1 and PMv (with exceptions in some temporal and prefrontal areas). Third, other than the exceptions noted above, there is a high degree of inverse de/resynchronization symmetry between the two hemispheres for both sensory and motor coding. Post-hoc analysis was conducted (below) to further examine these trends.

### Sensory and motor coding in specific regions of interest

Our next aim was to examine sensory and motor coding for specific regions of interest (Table 2), specific frequencies, i.e., α (7-15Hz) and β (15-35Hz), and through time. In order to simplify this and increase the power of our data, we followed a practice of previous MEG studies (e.g. (Van Der Werf et al. 2008)); supported by our observations of bilateral inverse symmetry (opposite power changed in left vs right hemispheres), we collapsed data across bilateral brain areas by subtracting data from corresponding left and right areas. Indeed, Supplementary Figures 2-16 show a strong left-right hemisphere symmetry for most areas. In the worst-case scenario, one hemisphere drives an effect alone (e.g. left M1 because of right hand use); however, subtracting the right hemisphere only adds noise but does not change our findings. Thus, to remain consistent in the summary analyses, we only show subtracted data in Figs. 6-7, but individual hemisphere data is provided for completeness in supplementary materials.

The steps in this analysis are illustrated in Figure 5. This uses the same example area as Figure 3 (mIPS) and begins where that analysis ends. Figure 5 shows sensory (upper row) and motor (lower row) power across multiple frequencies for left, right, and right-left mIPS respectively, followed by temporal plots of the α and β bands (left to right, all averaged across participants). As illustrated above, mIPS demonstrated very strong sensory coding. This is evident bilaterally in the strongly anti-symmetric synchronization and desynchronization in the first two panels, which results in a strong contralateral desynchronization in the following subtraction. The temporal plot shows this occurring in both the α and β bands at around 300 ms post-stimulus. Following the same sequence for the motor subtraction (lower row) reveals less power, but enough to yield bilateral contralateral synchronization in the α band, which peaks about 700ms before the movement. Analyses for all other brain areas considered are provided in Supplementary Figures 2-16. We followed the same procedure for all of the areas listed in Table 2, resulting in time courses for 16 bilateral cortical areas.

**Figure 5:**
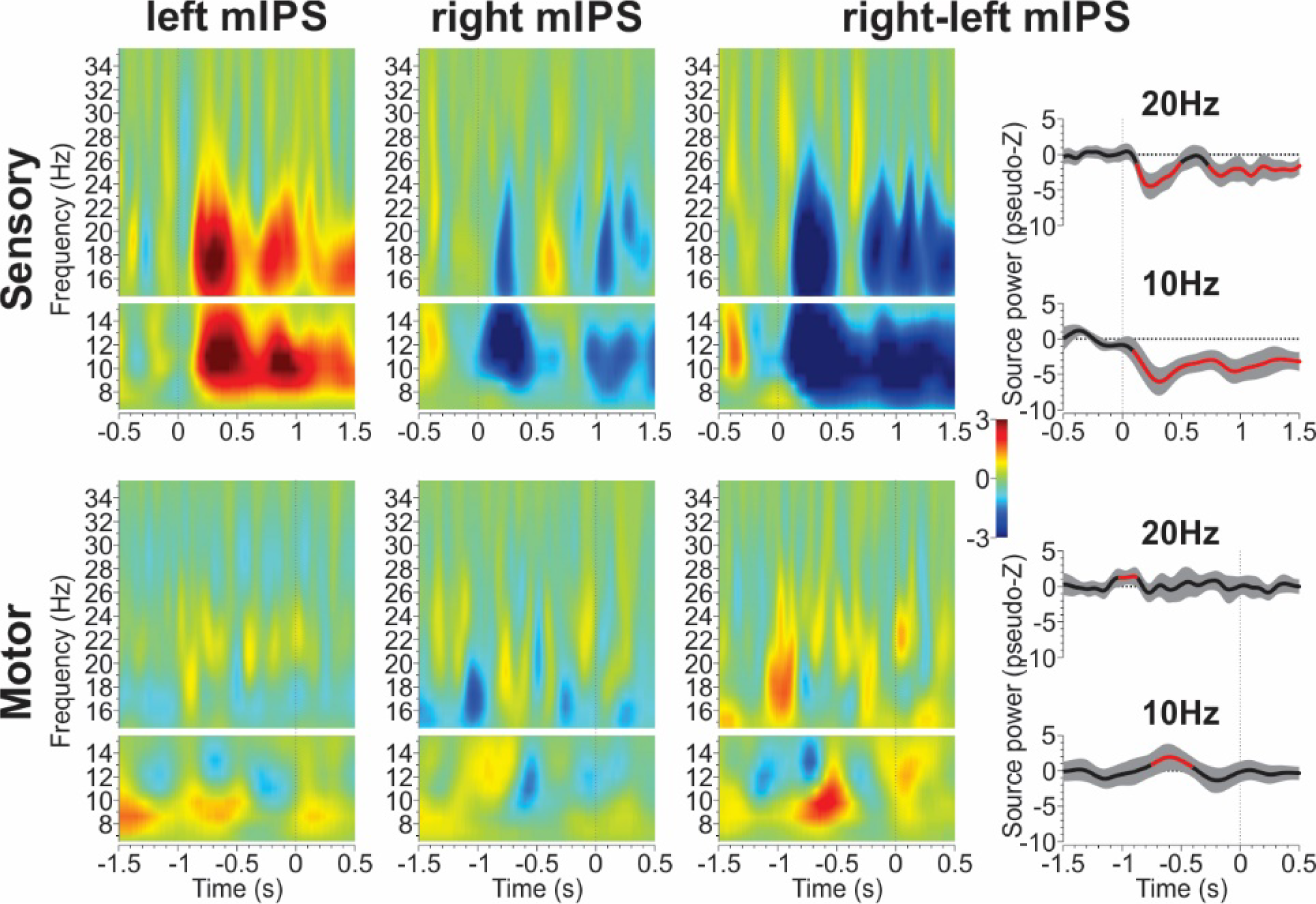
Time-frequency response (TFR) analysis of sensory-motor coding area mIPS. Top row shows sensory coding with cue alignment, bottom row shows motor coding with movement alignment. TFRs for left and right mIPS are shown separately in the first 2 columns. Red/resynchronization and blue/desynchronization with respect to baseline and with respect to left target/movement (due to left-right subtractions, see Methods). Taking advantage of the brain’s contra-lateral visual organization, we subtracted left from right TFRs in the third column to provide a single picture of activation. Since this is the result of L-R cue (movement) subtractions, red colors correspond to re-synchronization and blue correspond to de-synchronization of the brain area with respect to contra-lateral stimulus (movement) direction. Time course of α band power (10Hz) and β band poser (20Hz) is shown in the last column. Black curve and gray area indicate across participant mean and 95% confidence intervals. Red lines show activations that are significantly different from zero, i.e. different from baseline. mIPS showed strong sensory coding in α and β bands after cue onset and motor coding in α and β bands before movement onset.

To illustrate the results of this analysis, we plotted the average cue-related and movement-related whole-brain activations for each frequency band separately and added individual time courses of activation for each of our 16 bilateral regions of interest. These plots are shown in Figure 6 for the sensory code and Figure 7 for the motor code. For the whole brain analysis, the sensory code was computed during the 500ms post cue-onset (approximately the time window of peak sensory response) and the motor code was calculated during the 500ms before movement onset (approximately when motor planning responses peaked), with a mean temporal gap between these datasets of 815±83ms. Note that the whole-brain activations plotted on these figures show average power regardless of significance.

**Figure 6:**
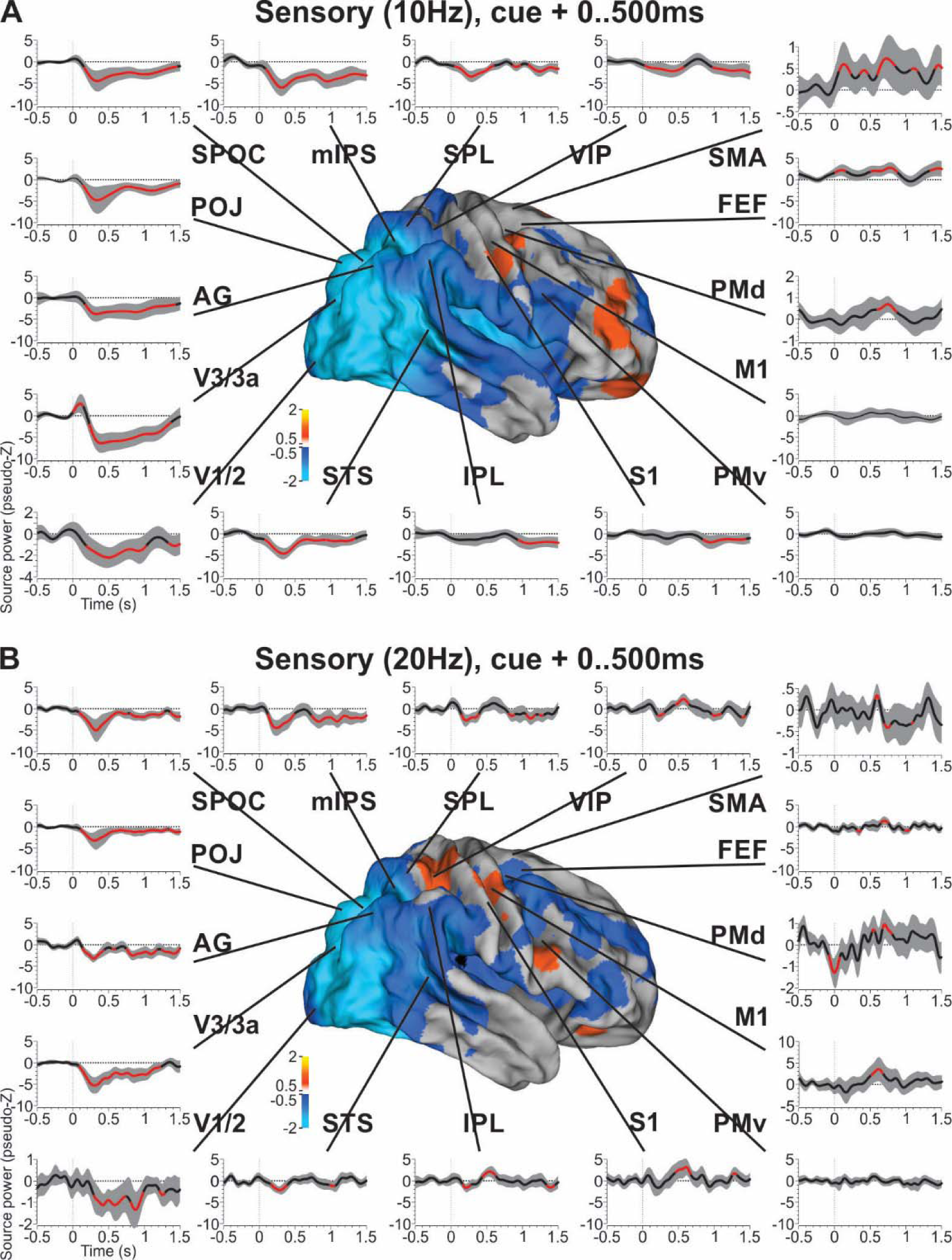
Summary of sensory coding across time in the whole brain. **A**. Whole-brain source power in the α band averaged across the first 500ms after cue onset and averaged across all participants. Individual time courses of sensory coding in the α band (10Hz) are shown for each brain area of interest. **B**. Same analysis for the β band (20Hz). Almost all areas (aside from PMv) showed significant changes in synchronization related to the sensory cue. Black curves and gray area indicate across participant mean and 95% confidence intervals. Red lines show activations that are significantly different from zero, i.e. different from baseline.

**Figure 7:**
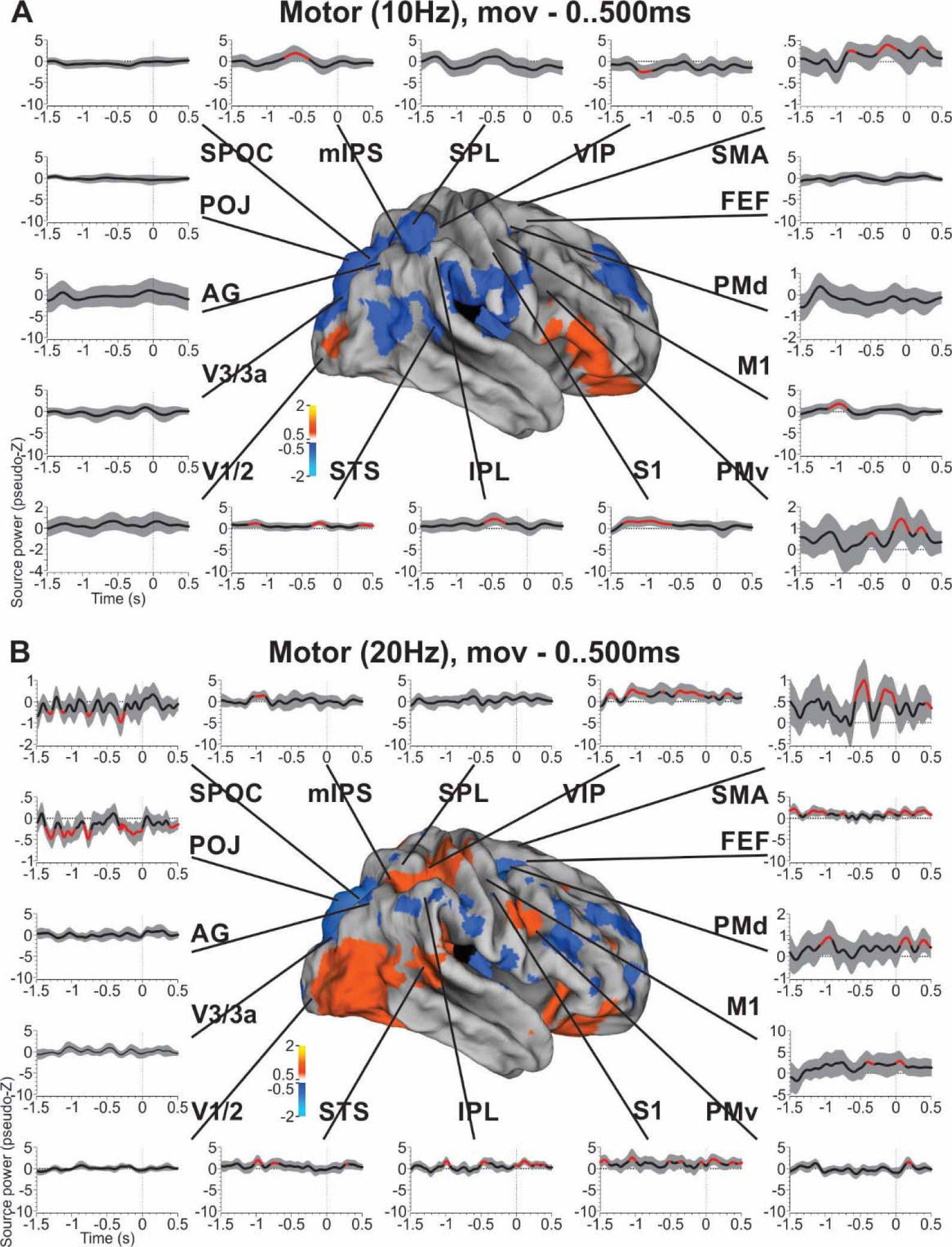
Summary of motor coding across time in the whole brain. Same conventions as in Fig. 6. **A**. Whole-brain source power in the α band (10Hz) was aligned to movement onset, averaged across the last 500ms prior to movement onset and average across all participants. Occipital areas did not show any motor coding. Significant motor codes in the α band appeared in parietal and frontal areas. **B**. B band (20Hz) motor coding was more prominent than α band motor coding, but only in parietal and frontal areas, not occipital areas.

As can be observed from Figure 6, most brain areas showed significant sensory coding during the delay period. This sensory response usually peaked 300 to 500ms after the cue onset, but sometimes persisted for more than 1 second. This was observed for both α (Fig. 6 A) and β (Fig. 6 B) bands and effects were strongest in occipital-parietal cortex, and tended to diminish along the posterior-to-anterior continuum of brain areas, with more frontal areas showing weaker and more variable sensory coding in both the α and β bands. Of interest is that while cue-related activation usually resulted in a desynchronization (relative to contralateral stimulation), some more traditionally motor-related areas (e.g. SMA, PMd, FEF) showed a cue-related re-synchronization of brain activity. It is also noteworthy that the initial sensory response, whenever present, appears to spread rapidly through the brain, almost appearing synchronously throughout occipital, parietal, and frontal cortex.

Inspecting motor-related activity in Figure 7, significant activations were observed in both α (Fig. 7 A) and β (Fig. 7 B) bands. Overall movement-related codes were less directionally selective and had more variable timing than the sensory code. Again, the α band signal was somewhat more stable through time, with more temporal variability in the β band. Earliest α and β motor activations occurred in STS, S1, VIP and M1, and were more prominent in motor-related areas (SMA, mIPS, PMv, IPL) closer to movement onset. However, overall the α band did not show as much significant motor-related activation as the β band. Indeed, we observed persistent β band motor activations (Fig. 7B) during most of the delay period in areas SPOC, POJ and VIP. Other movement-related areas exhibited motor activation closer to the movement onset, such as SMA and M1. Spatial lateralization of planning direction in S1/M1 might surprise neurophysiologists, but has also been observed in fMRI studies (Cappadocia et al. 2017).

### Sensory-motor transformation

As shown in Figures 5-7, many areas show both sensory and motor coding during the delay period of visual memory-guided wrist pointing. However, one cannot directly observe a transition between sensory and motor coding within and across areas from these separate sensory and motor analyses. To investigate this further, we computed a sensory-motor index (see Methods). This index captured the specificity of the coding scheme employed by an area on a millisecond basis, independently of frequency.

Results of this analysis are shown in Figure 8 and illustrate the gradual sensory-to-motor transformation across cortical space and time. As expected, we observed a series of areas showing only significant sensory codes, such as V1/2, V3/3a, SPL and FEF. For V1/2, V3/3a and SPL, strong sensory coding arose immediately after cue onset and was maintained for part of the delay period, but vanished prior to movement onset. This analysis also revealed areas that only showed significant coding for movement direction during the delay period, such as SMA, PMd and PMv. Those predominantly motor codes mostly emerged prior to movement onset toward the end of the delay period. Importantly, most brain areas in the identified network underlying the planning of goal-directed wrist movements exhibited early sensory coding followed by a progressive transition into motor coding. This was observed in areas SPOC, AG, POJ, mIPS, VIP, IPL, STS and M1. Interestingly, we observed a relatively clear early visual response in M1 and a rapid transition into reliable and significant motor coding (about 450ms into the delay period). Together with PMd, M1 showed the earliest significant motor code across all brain areas we investigated. These observations are further synthesized and summarized in the following section.

**Figure 8:**
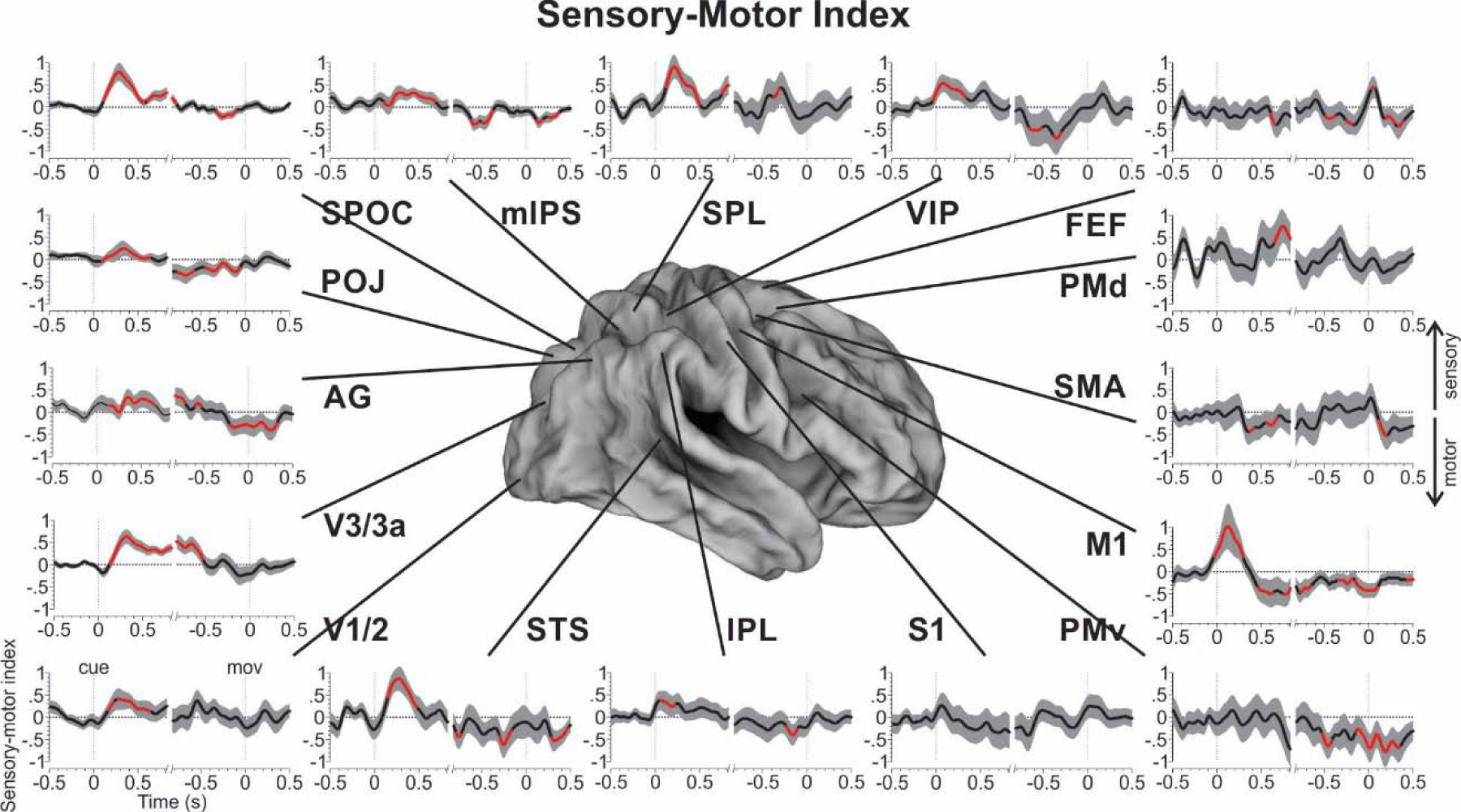
Sensory-motor index. The sensory-motor index (Eq. 3) is shown for each brain area as a function of time. Black curves and gray area indicate across participant mean and 95% confidence intervals. Red lines show indices that are significantly different from zero. The individual plots are split into cue alignment (-500..825ms around cue onset) and movement alignment (-825..500ms around movement onset) to account for variability in movement times (see Table 1). Index = +1 indicated perfect (noise-free) sensory coding; index = -1 indicates perfect motor coding. Index = 0 means that one cannot distinguish between sensory or motor coding.

### Summary: sensory and motor coding across cortical space and time

Figure 9 summarizes the data that we have described for sensory, motor, and sensorimotor coding in cue and premotor responses, and the relative timing of the sensorimotor transition between these sites, for the 16 bilateral cortical areas that we investigated in detail. In Figure 9A/B, the concentric circles placed at these sites represent whether sensory (cyan), motor (magenta), or neither (grey) coding is observed in (from center-out) the α band, β band, and sensorimotor index respectively. What this shows is an overwhelmingly uniform early sensory response to cue direction across occipital-parietal-frontal cortex (Fig. 9 A), with the exception of the sensorimotor index in PMd, and an overwhelmingly uniform movement direction response preceding the action in parieto-frontal cortex (Fig. 9 B), with the exception of the sensorimotor index in SPL.

**Figure 9:**
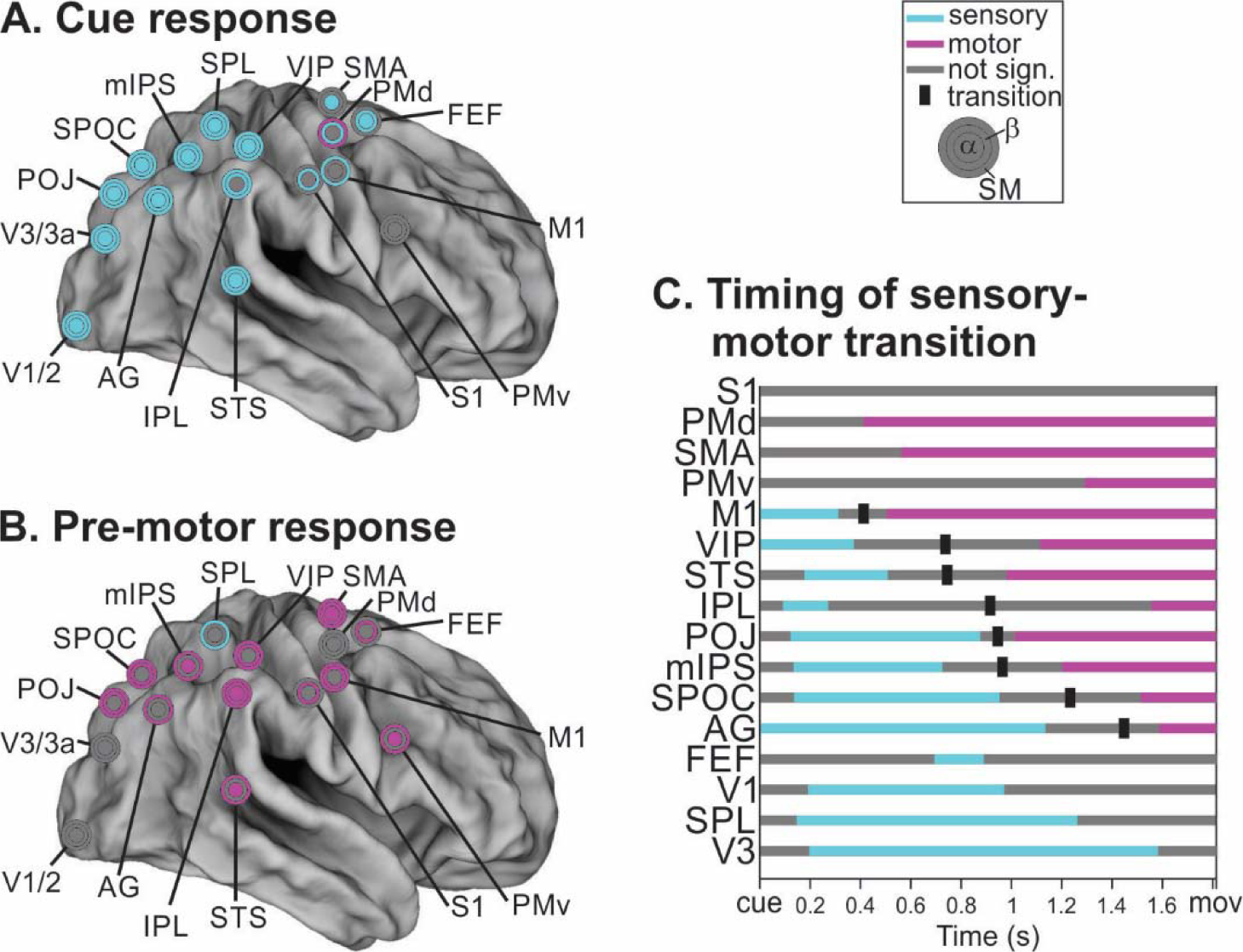
Sensory-motor transitions summary. **A**. Whole-brain view of coding schemes found in response to the cue within the first 500ms after the cue. Concentric disks indicate significant coding schemes for the α (inner disk) and β (middle disk) bands, as well as for the sensory-motor index (outer disk). Almost all cue responses show a sensory coding scheme across the brain. **B**. This coding scheme changes into a predominantly motor code during the pre-movement period (last 500ms prior to movement onset). **C**. Temporal evolution of coding schemes and timing of transitions across all brain areas according to the sensory-motor index. Earliest predominant motor codes appeared in PMd, SMA and M1 and the gradually appeared in more posterior areas, i.e. VIP, STS, IPL, POJ, mIPS, SPOC and AG (in order).

Importantly, Figure 9 C summarizes the temporal evolution of the sensorimotor transformation by showing the progression of the Sensorimotor Index through time. Again, cyan shows sensory coding, magenta shows motor coding, and the vertical ‘tick’ marks represent the cross-over point between significant sensory and motor coding (mid-way between last significant sensory and first significant motor index). These have been ordered, top-to-bottom, from earliest motor coding, to most persistent sensory coding. The striking result of this analysis is that, in response to a pro-anti instruction, a sensorimotor transition occurs over the course of approximately 1 second, and begins in frontal cortex, and then proceeds through parietal cortex toward occipital cortex, with clear transition points occurring mainly (but not exclusively) in posterior parietal cortex (AG, SPOC, mIPS, POJ, IPL, VIP)

## DISCUSSION

We set out to investigate where and when in the human brain visual sensory signals about a goal are transformed into appropriate motor commands. To do so, we took advantage of the natural dissociation of cue and movement directions in pro-/anti-tasks, and the high spatial-temporal resolution of MEG recordings across different frequency bandwidths. Contrasting our various conditions (Pro/Anti vs. Left-Right Targets) to each other in different ways elucidated sensory and motor coding-related activations. The results provide several important insights; first, if one takes snapshots in time, one observes predominantly sensory spatial coding throughout occipital-parietal-frontal cortex in response to a visual target stimulus; but prior to movement onset this switched to a predominantly motor spatial code in parietal-frontal areas. Interestingly, looking at the data in more detail, a progressive tendency from visual coding in more posterior areas toward movement coding in more frontal areas was evident. Further, a temporal sensorimotor progression could also be observed within most areas, especially in parietal cortex. Finally, in contrast to the very rapid (presumably forward) propagation of the sensory code, the sensorimotor response to the pro-anti cue was propagated backward from fontal cortex toward more posterior areas as time progressed. This is the first evidence showing the sensory-to-motor transformation in real time and at the whole-brain level in humans.

### Comparison to fMRI and Neurophysiology literature

The anti-saccade/reach paradigm has been used in conjunction with fMRI to study various aspects of motor suppression and preparation (e.g. (Connolly et al. 2000, 2002; DeSouza 2002; Curtis and Connolly 2007; Furlan et al. 2016)). The current results are most relevant to those studies which focused on the coding of visual vs. motor direction. In general, we were able to confirm that for some areas the directionality of sensorimotor activation (here in the form of cortical de-/re-synchronization) was primarily lateralized to the hemisphere contralateral to the visual stimulus and/or movement (Sereno et al. 2001; Medendorp et al. 2003; Medendorp 2004; Beurze et al. 2007, 2009, 2010; Fernandez-Ruiz et al. 2007; Bernier et al. 2012; Vesia and Crawford 2012; Chen et al. 2014). We also confirmed the general progression of spatial tuning for target responses in early occipital/parietal areas versus motor tuning in more parietal-frontal areas (Fernandez-Ruiz et al. 2007; Chen et al. 2014; Gertz and Fiehler 2015; Cappadocia et al. 2017; Gertz et al. 2017). Most relevantly, we confirm the general observation that motor directionality can be remapped within specific areas (Medendorp et al. 2003; Medendorp 2005), and more specifically the observation that anti-pointing can induce a sensory-to-motor transformation within and across many areas of parietal-frontal cortex (including SPOC, mIPS, and AG).

However, some of our detailed observations are harder to reconcile with the fMRI literature. For example, in the pro/anti wrist pointing task our MEG data seemed to be weighted more toward retrospective visual coding, whereas fMRI data were weighted more toward prospective movement planning signals (Gertz and Fiehler 2015; Cappadocia et al. 2017; Gertz et al. 2017). This apparent discrepancy was in fact due to the left-right target / pro-anti condition / left-right hemisphere contrast; indeed, average motor activity was generally very strong, especially in M1 (data not shown), but here we only focused on the spatial/condition contrasts. Furthermore, fMRI experiments consistently show more activation in the hemisphere contralateral to the effector (Medendorp 2004; Gertz and Fiehler 2015; Cappadocia et al. 2017), whereas we did not observe this in our contrasted MEG data; if anything, ipsilateral activity was often greater, with the exception of S1 and M1. Also, the sensory-motor remapping that we observed here was even more widespread in our recent fMRI experiment (Cappadocia et al. 2017), for example extending to occipital cortex and premotor cortex. This might have something to do with fMRI’s relatively greater sensitivity to input to areas (Logothetis 2008), MEG’s insensitivity to gyri, or our focus here on the α and β bands. In general, we do not take these as contradictions, but rather as complementary findings that likely reveal technical limitations in these different approaches.

At the microscopic level, many neurophysiological studies have demonstrated task-induced remapping of visual information (e.g., (Duhamel et al. 1992; Dash et al. 2015)). Specifically, anti-saccade/reaches induce directional remapping within areas and even in specific parietal cells (Matthews et al. 2002; Zhang and Barash 2004; Gail et al. 2009). Such responses likely underlie the cue-dependent transformations derived from MEG (Van Der Werf et al. 2008), and here we extend this to a much broader network for goal-directed movement planning. Finally, it has also been demonstrated in the gaze control system that target coding transitions to motor coding across specific cell types in the superior colliculus, parietal cortex, and frontal eye field during memory-guided pro-movements (Sadeh et al. 2015; Sajad et al. 2015, 2016), suggesting that many of the observations gained from the pro/anti task may generalize to everyday movements.

### Frequency-Dependence

A major advantage of current MEG methodologies over any one of fMRI, unit recording, or EEG is the ability to dissect the power of oscillations across various frequency bandwidths from the entire cortex, and localize these oscillations to specific brain sites (Alikhanian et al. 2013; Cheyne 2013). As noted in the methods, we focused on the α and β bands because the literature suggests these are most closely linked to sensorimotor events, but similar observations were made in the γ band during anti-remapping of saccade targets (Van Der Werf et al. 2008). The prominence of both sensory and motor signals in the α-band in our study, and its role in sensorimotor transformations are consistent with the ability of α-band TMS over parietal cortex to disrupt both target memory (SPOC) and reach vector planning (mIPS, AG) (Vesia et al. 2010; Vesia and Crawford 2012).

While the α band more strongly reflected sensory processing, motor planning seemed to invoke β band oscillation changes. While this was not surprising, we did not expect sensory processes to modulate β power and motor processes modulate α power to the extent they did. However, different frequencies of oscillation are believed to arise from recurrent processing with different loop delays (α band through cortical-thalamic interactions (Suffczynski et al. 2001); β band through cortico-cortical coupling (Cabral et al. 2014)). We believe that the lack of frequency specificity of sensory-motor processes might reflect the involvement of complex networks relying on more or less sub-cortical processing rather than being of any direct functional significance with respect to the sensory-motor task at hand.

### Timing

The other fundamental advantage of MEG over fMRI in investigating source-localized activation is real-time measurements. Given that fMRI typically has a temporal resolution of around 2 seconds (theoretically as low as 100ms in fast-event designs), it cannot possibly match the resolution of MEG, and certainly has not done so in this specific area of research. Our results suggest that the response to the visual stimulus propagates rapidly through the occipital-parietal-frontal axis, in agreement with numerous neurophysiological studies (see above). Presumably this reflects a normal process that would also initiate movement coding in pro-movement reaction-time tasks. However, the pro-anti task introduces an additional top-down transformation. Here, we were able to show that in the case of our task, this transformation was initiated in frontal cortex, and then spread progressively backwards, presumably through recurrent connections, through more posterior regions over the course of a second. This is consistent with neurophysiological studies which found earlier activation of frontal over parietal cortex in certain tasks (Schmolesky et al. 1998; Omrani et al. 2016). Re-activation of occipital areas has also previously been reported in the grasping system (Monaco et al. 2017). Finally, this finding could explain the classic observation that frontal cortex damage specifically impedes function in anti vs. pro-movement tasks (Guitton et al. 1985).

Interestingly, this view of parietal cortex representing the current state of affairs instead of performing actual computations is consistent with recent proposals both from the motor control and the decision-making communities. Indeed, there is strong evidence for posterior parietal cortex acting as a state estimator for motor control (Ogawa et al. 2007; Mulliken et al. 2008; Shadmehr and Krakauer 2008; Andersen and Cui 2009; Grafton 2010; Shi and Buneo 2011; Marigold and Drew 2017). Similarly, it has recently been suggested that decision states might only be conveyed to parietal cortex after the decision outcome has been computed elsewhere (Latimer et al. 2013; Katz et al. 2016; Huk et al. 2017). In our data, the motor intention in parietal cortex was updated after frontal areas integrated pro-/anti-instructions or task demands (DeSouza 2002; Everling and DeSouza 2005), thus reflecting (but perhaps not actively computing) the current intention. We believe that this is an intriguing hypothesis that should be examined in future studies.

### Implications for Models of sensory-motor transformations

Two types of conceptual models have been proposed regarding the way the brain could compute the sensory-to-motor transformation. The most popular class of models uses artificial feed-forward neural networks and suggests that visuomotor transformations occur serially through successive stages of processing across different brain areas (Zipser and Andersen 1988; Pouget, Deneve, et al. 2002; Pouget, Ducom, et al. 2002; Blohm et al. 2009; Blohm 2012). This model class predicts that the inherent reference frame of coding within a brain area is fixed across time. Alternative models take advantage of the dynamic nature of brain signals and suggest that sensorimotor transformations can be carried out over time within a single area receiving all relevant inputs (Deneve et al. 2007; Keith et al. 2010; Schneegans and Sch??ner 2012). If this was true, we would expect the spatial coding scheme within a given brain area to change over time.

Our MEG results suggest that both models are incomplete and require revision. Indeed, our data suggest that sensory-motor transformations occur simultaneously both across space and across time (Sajad et al. 2016). In addition, the exact temporal transition does not seem to align with the spatial gradient, i.e. premotor and motor cortex are at the motor coding end of the spatial gradient, but the motor code emerges earliest in those areas over time (see Discussion below). It is unclear what the reason for this apparent contradiction is. It is also unclear why so many areas are involved in the sensory-to-motor transformation. It can only be speculated that the reason for the latter might lie in other factors of the sensory-to-motor transformation that were not considered in this study, i.e. effector choice, posture integration, reference frame transformations or target selection / decision making processes. Overall, our findings call for a new dynamic model of sensory-to-motor transformations for goal-directed movements.

M1 and PMd showed the earliest motor codes. The fact that motor coding in other sensory-motor areas occurred later, could have two distinct reasons: (1) since the sensory-motor index captures predominant coding schemes, earlier motor codes could be masked by stronger sensory coding, but both could co-exist; (2) the sensory-to-motor transformation first occurs in a feed-forward fashion from occipital to frontal areas and then feedback connections gradually update earlier areas to reflect the upcoming motor plan as represented in frontal cortex. These hypotheses would be best dissociated in future non-human primate electrophysiology studies. If the latter turned out to be true, then a dynamic bi-directional hierarchical model – such as Tsotsos’ selective tuning model for attention (Tsotsos et al. 1995; Tsotsos and Kruijne 2014) would be best suited to describe sensory-motor planning processes in the brain. Such a bi-directional model would also be in line with recent suggestions of learning hierarchies in the brain (Roelfsema and Holtmaat 2018), which would provide a fascinating new perspective – state estimation in parietal cortex driving sensory-motor learning.

### Limitations of MEG and the current study

Our findings are likely incomplete due to measurement limitations of MEG. Indeed, in theory MEG recordings are most sensitive for brain areas in the wall of sulci, i.e. when cortical columns are parallel to the scalp surface. Ideally, to overcome this limitation complementary EEG signals should also be recorded and analyzed in conjunction with MEG signals. Practically however, these limitations are less severe for 2 reasons. (1) The gyral regions to which MEG sensors are theoretically insensitive are extremely small (Hillebrand and Barnes 2002; Goldenholz et al. 2009) and typically well within the extent of MEG current spread. (2) Few brain areas are strictly orthogonal to the scalp as the extent of brain areas usually involves some non-orthogonal regions. Thus, while MEG is less sensitive to gyral regions, this is less of a concern in practice (Hillebrand and Barnes 2002; Koser 2010; Cheyne 2013; Baillet 2017).

Another limitation is the spatial resolution of MEG. This could in particular be an issue for regions that are spatially very close to one another. In that case one might ask if different areas can be meaningfully resolved. While we cannot be completely certain that this was the case, we do show that results can be quite different for areas that are spatially close together. This can be observed when comparing results for FEF, PMd and SMA. While we cannot rule out “current spread”, these observations make us confident in our findings.

To obtain population significance values, we averaged our data across participants and across trials. This averaging process could have smeared out single-trial and/or single-participant transformation dynamics / timing. However, we still found distinctive time courses between areas. For example, motor codes emerged within about 450ms in PMd and M1, whereas it took over 1s in other areas such as PMv for example. Therefore, we think that even if temporal smearing did occur, our task was still able to reveal timing differences between areas. This is interesting because it means that different parts of the network seem to carry out the sensory-motor transformation at different points in time, or at least they reflect sensory vs motor codes at different points in time during the delay period.

Our contrast / subtraction approach to reveal sensory or motor coding relies on spatial selectivity with brain areas as well as an assumption of linearity of effects. Spatial selectivity is present in many brain areas. However, that does not necessarily imply that the net magnetic signal is affected differentially by our conditions. In other words, the neural population code might undergo task-related changes that do not result in a net MEG signal change. As a consequence, we might underestimate the number of areas involved in sensory-motor transformations for goal-directed movements, and / or we might not be able to detect changes in coding within an area. The assumption of linearity could also lead to shortcomings in interpreting MEG signals. Here, linearity designates the symmetry in oscillatory changes with identical but opposite stimulus / movement changes. If such linearity was not given, then our approach of computing sensory/motor indices might either miss certain effects or erroneously find effects (false positives). While it is possible that nonlinear effects exist in our data, the signal-to-noise ratio in our experiment was too low to identify them. The robustness of results across participants (as revealed by our statistics) indicates that the linear assumption leads to an acceptable first approximation.

Sensory-motor transformations are composed of many conceptual steps, including target selection, reference frame transformations, effector selection and accounting for arm posture. As a starting point of whole-brain MEG analyses of the goal-directed movement network, we only address how sensory signals of target location are converted into appropriate motor commands. Other studies are required to inspect other aspects of sensory-to-motor transformations, such as the influence of effector choice and posture (Kakei et al. 1999, 2001; Beurze et al. 2009; Leone et al. 2014; Heed et al. 2016; Fujiwara et al. 2017).

## Conclusions

Planning a movement requires the conversion of visual information into a goal. Our whole-brain MEG analysis has uncovered several novel findings: (1) the initial occipital-parietal-frontal sweep of sensory information was followed immediately by the appearance of a motor code resulting from processing of the pro-/anti-cue information. (2) This motor code appeared first in traditional motor areas (M1, PMd) within 500ms of cue presentation. (3) Motor coding then spread gradually to more posterior areas over time, as if parietal cortex received an update of the motor intention from motor areas.

## Acknowledgments

Experiments and H.A. were supported by a Canadian Institutes for Health Research Grant held by JDC. GB was supported by a Marie Curie Fellowship (EU) during the experiments and by NSERC (Canada) thereafter. JDC is supported by a Canada Research Chair. The authors would like to thank Andrea Boston and Sonja Bells for technical assistance and Dr. Paul Ferrari for comments on the current manuscript.

